# An evolutionary algorithm for designing microbial communities via environmental modification

**DOI:** 10.1101/2020.11.25.398644

**Authors:** Alan R. Pacheco, Daniel Segrè

## Abstract

Despite a growing understanding of how environmental composition affects microbial community properties, it remains difficult to apply this knowledge to the rational design of synthetic multispecies consortia. This is because natural microbial communities can harbor thousands of different organisms and environmental substrates, making up a vast combinatorial space that precludes exhaustive experimental testing and computational prediction. Here, we present a method based on the combination of machine learning and dynamic flux balance analysis (dFBA) that selects optimal environmental compositions to produce target community phenotypes. In this framework, dFBA is used to model the growth of a community in candidate environments. A genetic algorithm is then used to evaluate the behavior of the community relative to a target phenotype, and subsequently adjust the environment to allow the organisms to more closely approach this target. We apply this iterative process to *in silico* communities of varying sizes, showing how it can rapidly identify environments that yield desired phenotypes. Moreover, this novel combination of approaches produces testable predictions for the *in vivo* assembly of microbial communities with specific properties, and can facilitate rational environmental design processes for complex microbiomes.

## Introduction

Microbial communities are complex ecosystems that are crucial to the health and function of many biomes, from aquatic ecosystems to the human gut [1–5]. In addition to yielding a growing understanding of the composition of various microbial ecosystems [6–8], recent advances in DNA sequencing and synthetic biology have enabled new efforts to engineer synthetic multispecies consortia for a variety of applications [9–11]. For example, multispecies systems have been designed to degrade complex substrates or pollutants [12–15], as well as to produce biofuels and molecules for human consumption [15–18]. Advances such as these portend the advent of new applications in synthetic ecology, in which communities of microbes can be readily designed for a vast number of useful outputs. However, this promise is hampered by the difficulty in genetically manipulating individual organisms at community scales, as well as by the lack of a mechanistic understanding of how environmental factors and interspecies interactions shape communities [19–21]. These challenges open up the important question of whether a more accessible parameter, i.e. the chemical composition of the environment, can be modulated to confer specific functions on microbial consortia.

A number of studies have demonstrated the crucial role that changes in environmental composition play in defining microbial community phenotypes, such as in the gut microbiota [22,23] and in aquatic and terrestrial ecosystems [24,25]. As natural ecosystems contain complex combinations of different nutrients, studies have also begun to disentangle the nonintuitive relationship between community properties and resource identity and heterogeneity [24,26–29]. These observations point to the manipulation of environmental composition as a promising method for producing synthetic consortia with defined functions. However, these and other recent studies have demonstrated that community growth and structure can be so sensitive to environmental composition that even closely-related environments can produce very different communities [28,30]. Therefore, in order to reach a phenotype of interest, in practice it often remains necessary to explicitly test a multitude of different specific nutrient combinations – a task that can quickly become experimentally intractable. For example, screening a consortium under all combinations of 20 nutrients – a quantity vastly lower than the number of unique metabolites found in natural settings – would require 1.05 million individual experiments, a scale that remains inaccessible to current conventional microbiological methods. Organism-specific computational models can be deployed to run *in silico* analogs of these experiments [31–33], though the number of simulations required would also rapidly become computationally intractable for more complex environmental search spaces.

To begin addressing these challenges, we present here the design of a genetic algorithm (GA) framework to rapidly identify environmental compositions that result in target community phenotypes. Our method, conceptually similar to processes used to evolve communities toward specific functions [34–36], searches large spaces of nutrient combinations to produce candidate environmental compositions that optimize a given ecological objective. To test our GA framework, we rely on a large set of *in silico* community experiments consisting of over 6,000 unique environment-community pairings. While ultra-high-throughput experimental platforms can also generate large sets of community phenotypes in combinatorial environments [37–39], we employ a purely computational technique to demonstrate the ability of our genetic algorithm to identify environments that confer specific target community taxonomic compositions and desired patterns of metabolic secretion and exchange. In addition, we show how this pairing of an evolutionary algorithm with models of community ecology allows us to maximize community objectives under a much larger (~600,000 environments) combinatorial space. In sum, this method is able to reach a desired community design goal without requiring explicit ecological parameters as input, allowing it to serve as a versatile tool for exploration of large combinatorial spaces and future applications in experimental synthetic ecology.

## Results

### Generation of microbial community phenotypes in combinatorial environments

In order to test our search algorithm, we first simulated the growth of multispecies microbial communities under a large number of environmental compositions. This was done via a dynamic flux balance analysis (dFBA) technique [40] using the COMETS software package [41,42], which enables a mechanistic evaluation of community growth and metabolic exchange using experimentally-validated computational models of individual organisms (see Methods). Predictions using dFBA have been shown to recapitulate key microbial phenotypes, while also generating broader statistical mappings of community structure and interactions [32,33,43,44]. Moreover, the use of these models allows us to enumerate a complete set of environment-phenotype mappings that is large yet computationally tractable, allowing us to identify every possible community outcome and evaluate the quality of solutions identified by our algorithm against a known optimum. Our mapping was generated by simulating the growth of 13-species communities in a variety of environmental compositions. The *in silico* organisms that make up our communities were selected as they represent a diverse cross-section of taxa and metabolic capabilities (see Methods), in theory allowing us to maximize the variability of yields, taxonomic compositions, and interspecies interactions across different environments. We used combinations of up to 4 of 20 different carbon sources in order to generate a total of 6,196 unique environmental compositions, to which we added equal amounts of all 13 organisms (see Methods). The growth of these communities in all environments was then simulated with COMETS over a 24-hour timespan.

Our simulated communities displayed high degrees of compositional variability across the environmental conditions we tested (Figure 1a). Specifically, six *in silico* organisms (*B. subtilis, E. coli, P. aeruginosa, S. boydii, S. coelicolor*, and *S. oneidensis*) reached relative abundances of more than 50% in at least one environment, and all organisms encountered at least one environment in which they could not grow. Organism relative abundances displayed mean variances of 0.02, mean species richness values of 3.30 ± 0.99, and mean Shannon entropy values of 1.29 ± 0.49 (Figure 1b, c), which were comparable to those of similarly-sized communities assayed experimentally [28]. Moreover, we encountered a wide distribution in the number of metabolic exchanges across environments. We define a metabolic exchange as the transfer of a unique metabolite from one organism to another, and found that our environments resulted in 435.49 ± 106.49 such transfers on average (Figure 1d). Interestingly, the failure of six organisms (*K. pneumoniae*, *L. lactis*, *P. gingivalis*, *R. sphaeroides*, *S. cerevisiae*, and *Z. mobilis*), many of which have a variety of metabolic auxotrophies [45–48], to grow in any environment suggests that the levels of metabolic exchange observed in our communities was not enough to sustain these more specialized organisms. In sum, these results recapitulate elements of the often nonintuitive relationship between environmental composition and community structure observed in nature, serving as a suitable dataset on which to test our search algorithm.

**Figure 1.**
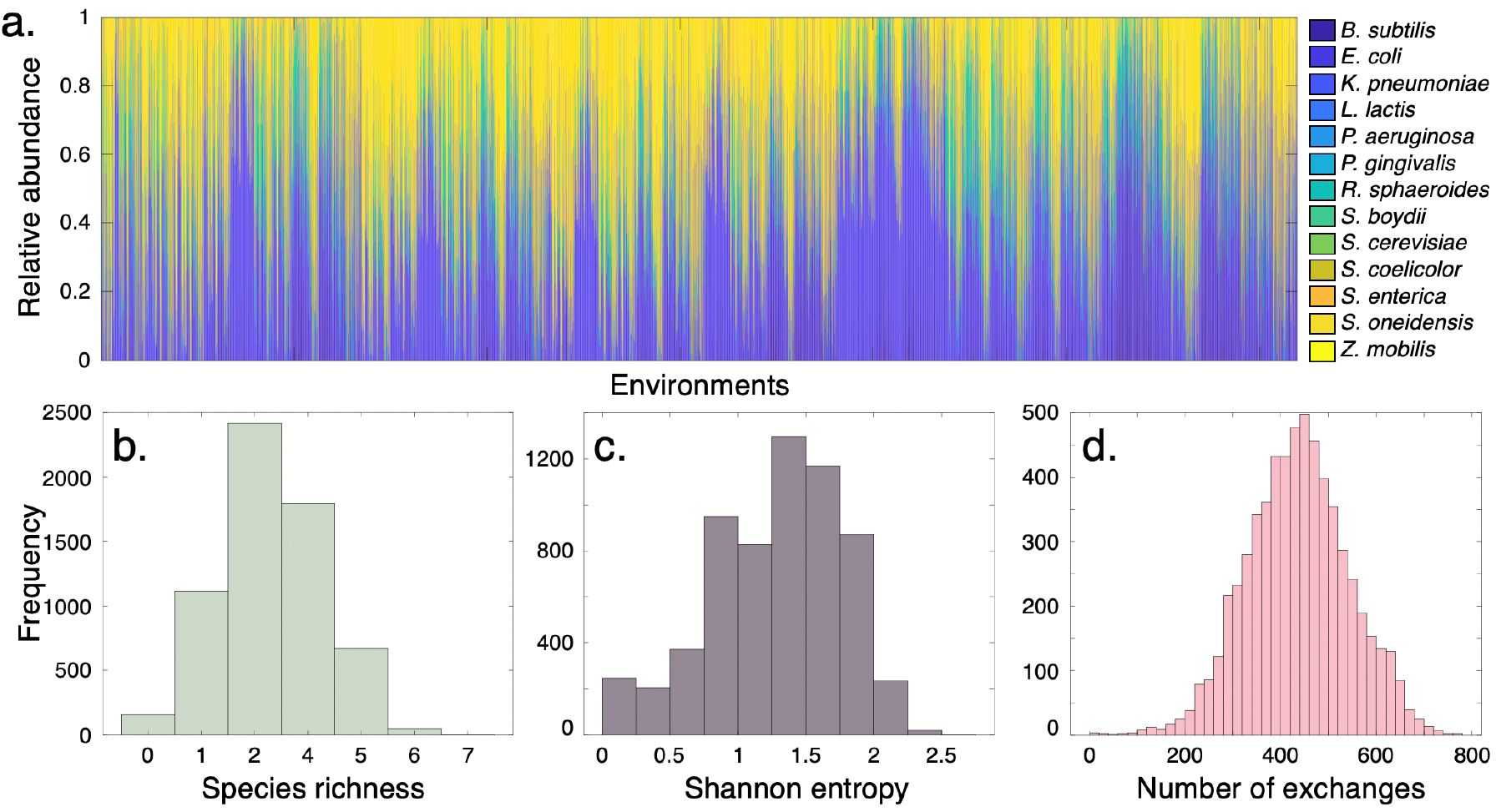
Structural and ecological properties of simulated 13-species communities. **(a).** Relative abundances of organisms after 24 hours of growth in all 6,196 combinatorial environmental compositions. **(b-d).** Distributions of species richness **(b)**, Shannon entropy **(c)** and total number of exchanges **(d)** observed across all environments. Here, one exchange is defined as the transfer of a unique metabolite from one organism to another, e.g. the secretion of metabolite *m* by organism *A* and its absorption by organism *B* represents one exchange. As our simulations contained 737 unique metabolites, the total possible number of exchanges (i.e. if each organism transfers each metabolite to each other organism) totals 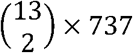, or 57,486.

### A simple evolutionary algorithm rapidly identifies environmental compositions

Having generated a wide array of phenotypic data, we designed a search algorithm to identify environments within our dataset that would result in specific community properties. This method, a genetic algorithm based on the process of natural selection [49–52], functions as follows: first, a population of *P* environmental compositions is generated, each containing a random assortment of a maximum of *N* unique nutrients. Community phenotypes (e.g. species abundances, interspecies interactions, metabolic secretions) on each environment are recorded, and each environment is scored according to the community function being optimized. A subset *σ* consisting of the top-performing environments is then selected to be propagated to the next generation. The remaining *P* – *σ* environments are generated by combining nutrients contained in the top *σ* environments (crossover), and by introducing new nutrients (mutation) at rates defined by a parameter grid search (Supplementary Figure 1). These new *P* environments are provided to the *in silico* communities, and the optimization process continues for *G* generations (Figure 2). The objective of the algorithm is therefore to converge to a final set of environmental compositions that confer the desired properties on the community being tested, without prior knowledge of the dataset’s structure.

**Figure 2.**
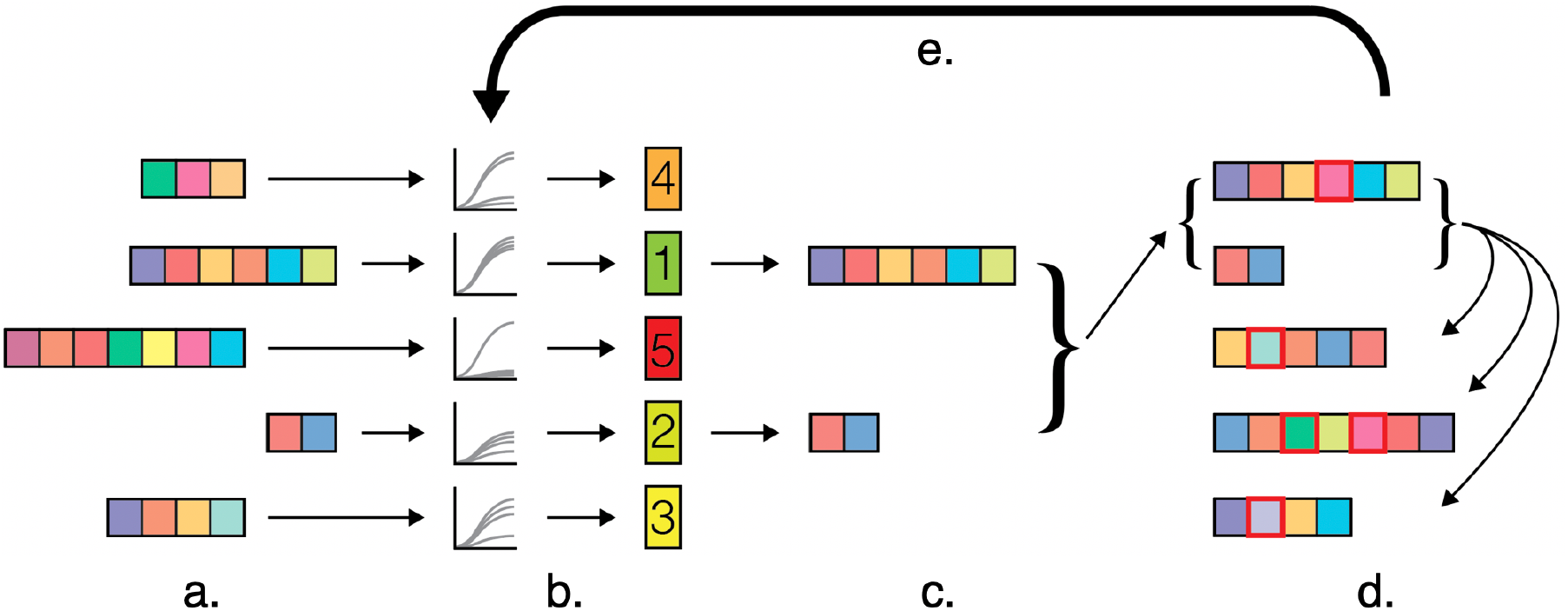
Schematic of genetic algorithm process for microbial community design. (*a).* A set of *P* environmental compositions, each containing a varying number of limiting nutrients, is randomly generated. (*b*). The community phenotype observed in each environment is determined. As a representative example, this figure shows the GA process with taxonomic balance as the objective to be optimized. The environments are ranked according to their resulting communities’ taxonomic balance, and (*c*) the top *σ* environments are selected. Here, the environments that yielded the top *σ* = 2 taxonomically balanced communities are chosen. (*d).* A new population of *P* environments is generated. First, the top *σ* environments are carried over into the new population as ‘parents’, and the remaining *P* – *σ* ‘offspring’ environments are generated via multipoint crossover (i.e. the individual nutrients in the parents are shuffled to produce heterogeneous offspring). Variation is introduced into the new population via mutation, in which each individual element has a defined probability of being changed into a new one (red squares). (*e*). The process of environment ranking, propagation, crossover, and mutation is carried out for a total of *G* generations.

We first applied this framework to identify environments that would maximize the final taxonomic balance of our communities. Though it is uncommon for organisms to be equally represented in natural settings [53–58], coexistence of multiple organisms is a desirable property for engineered consortia as it can enable tasks useful in biotechnology, such as metabolic division of labor [19,59]. As such, we sought to identify environments that resulted in relatively even species abundances. To do this, we applied the genetic algorithm to search for environments that would maximize the Shannon entropy of our *in silico* communities (see Methods). In order to gain a statistical representation of its performance, we ran our algorithm 50 separate times, each with different random initial compositions of *P* = 10 environments. For each GA process, we recorded the generation at which the algorithm’s proposed solutions crossed the 99^th^ percentile of all solutions as a way to quantify its performance. We found that, on average, our algorithm identified solutions that exceeded the 99^th^ percentile of Shannon entropy values after approximately 3 generations (Figure 3a, Supplementary Table 3). As each generation tested *P* = 10 environments, this performance represents explicitly testing only 30 unique *in silico* experiments out of a total of 6,196 total nutrient combinations. Though the algorithm generally converged quickly to near-optimal solutions, we observed some variability in the specific environmental compositions it selected. For this particular objective, our method resulted in 12 distinct environmental compositions across the 50 different random seed environments, all of which showed high degrees of consistency and taxonomic balance in the resultant communities (Figure 3a, inset).

**Figure 3.**
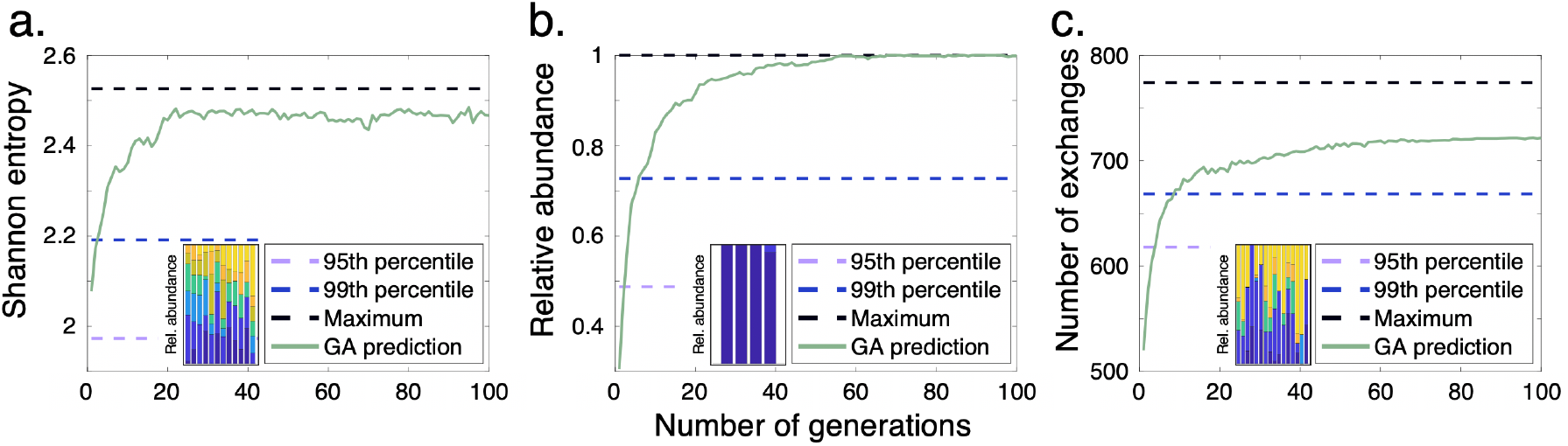
Performance of genetic algorithm on various ecological objectives. The average number of generations (using 50 random seed sets of *P =* 10 environments) required to identify environments that surpassed the 99^th^ percentiles of (*a*) community Shannon entropy, (*b*) the relative abundance of *B. subtilis*, and (*c*) the total number of metabolic exchanges between organisms. Insets show the organism relative abundances of the top environmental conditions identified. All quantities, including results for optimization of the remaining 12 organisms’ relative abundances, are found in Supplementary Table 3.

In addition to optimizing general ecological properties, we tested the capability of our algorithm to identify environments that would maximize more specific features. We first chose to optimize the relative abundances of individual organisms and selected *B. subtilis*, which grew in 2,130 out of 6,196 environments (Figure 1a), as a representative example. Again using 50 random seed sets, we found that the genetic algorithm was able to identify solutions that exceeded the 99^th^ percentile of *B. subtilis* abundances after approximately 8 generations on average (Figure 3b). We found that our algorithm converged to fewer distinct environmental compositions for this objective across our 50 random seeds, from which 4 distinct environments emerged (Figure 3b, inset). These environments nonetheless all conferred very high relative abundances to *B. subtilis*. Testing this capability for the remaining organisms revealed that similarly low numbers of generations were required to converge to optimal solutions (Supplementary Table 3), demonstrating the utility of this framework to identify environments that maximize individual species abundances.

Interspecies metabolic cooperation, often associated with microbial ecosystem stability, is a common target mechanism for community engineering [60,61]. Nonetheless, identifying environments that lead to the emergence of specific interactions remains an elusive task. We thus sought to determine whether our GA framework could also identify desired patterns of metabolic exchange from our computational dataset. We set the total number of interspecies exchanges as our objective function, in order to identify the environments that would maximize metabolic cooperation across all organisms. Our genetic algorithm was able to identify environments that surpassed the 99^th^ percentile of metabolic exchanges after 6 generations on average, representing a total of 60 *in silico* experiments (Figure 3c, Supplementary Table 3). Notably, the selected environments resulted in varied taxonomic compositions, ranging from those with high abundances of *E. coli* and *S. oneidensis* to those with more balanced compositions (Figure 3c, inset). This result suggests that the degree of metabolic exchange does not necessarily correlate with community taxonomic composition in our dataset, which parallels experimental observations showing conflicting correspondence between taxonomic structure and ecological function [62,63].

Given its ability to optimize the general prevalence of interspecies interactions, we also tested our algorithm on more specific patterns of secretion and exchange. In particular, we sought to determine if we could identify environments that resulted either in greater metabolic flux toward one particular organism or in greater overall secretion of a particular metabolite, as such specific phenomena are commonly leveraged for synthetic community design [60,61]. We again used *B. subtilis* as a representative organism to test the former capability, finding that our GA identified environments that surpassed the 99^th^ percentile of metabolic exchanges toward this organism after 9 generations on average. Testing the same capability with our remaining organisms as targets showed similar performance (Supplementary Table 3). We next set the net community-level output of specific metabolites from all organisms as an optimization target, in order to identify environments that would maximize their secretion. To do this, we selected 24 metabolites: 12 that were most highly secreted across all 6,196 simulations, and 12 that were least secreted. For the former set, we found that our algorithm identified solutions surpassing the 99^th^ percentile of secretion after, on average, 11 generations (Supplementary Table 3). However, our algorithm’s performance suffered for metabolites with low secretion flux, requiring on average 143 generations to reach this same benchmark.

Despite eventually converging to near-optimal solutions for all of the metabolite secretion patterns we tested, the longer convergence time needed to identify solutions for some metabolites prompted us to quantify its dependence on the number of times a particular metabolite was observed to be secreted across all simulations. We thus analyzed the average number of generations needed to surpass the 99^th^ percentile for a given target metabolite with respect to the number of times it was observed in our dataset, finding that these two quantities were inversely proportional to each other (Supplementary Figure 2a). Though this effect reveals a limitation of our method (or indeed of FBA itself), a large number of generations is needed for a rare minority of objectives. For this dataset, we determined that the secretion of 61.4% of organic metabolites could be maximized within 50 generations, with only 21.5% of metabolites requiring over 100 generations (Supplementary Figure 2b).

### Searching for community phenotypes in larger combinatorial spaces

Having benchmarked our GA framework on an exhaustive environment-phenotype mapping, we aimed to test its performance in a much larger search space. We thus applied it to determine whether certain environmental compositions could yield communities with highly specific organism relative abundances. This goal draws from efforts to precisely control organism ratios in mixed cultures, which is particularly relevant for synthetic communities applied to the synthesis of biofuels or chemicals [64–66]. Here, we sought to identify environments that would allow one of three organisms – *B. subtilis*, *E. coli*, and *S. coelicolor* – to reach a high abundance in a community (95%), while maintaining the remaining two at low abundances (2.5% each). We used a list of 154 limiting carbon sources from which we allowed our algorithm to select a maximum of 3. This search space, consisting of 596,904 unique environmental compositions, remains computationally expensive to test exhaustively using ecological modeling methods like dFBA and nearly impossible to test experimentally. Therefore, this application illustrates the capability of our GA framework to operate in an exploratory fashion within spaces that cannot be fully mapped.

To search this larger combinatorial space, we carried out dFBA simulations of our community in the selected environments as they were produced by the genetic algorithm, instead of generating a full environment-phenotype mapping *a priori* as above (see Methods). The quality of these environments was assessed by calculating the sum squared error (SSE) between the resulting community compositions and our target abundances [0.95, 0.025, 0.025], and the objective of the GA was therefore to minimize this quantity. We found that, by iteratively searching this large combinatorial space, the GA framework successfully identified environments that allowed each organism to reach a high relative abundance while keeping the remaining two at low, but nonzero abundances (Figure 4a-c). Notably, the algorithm converged on multiple such environmental compositions, indicating a type of metabolic flexibility with regards to specific final taxonomic compositions.

**Figure 4.**
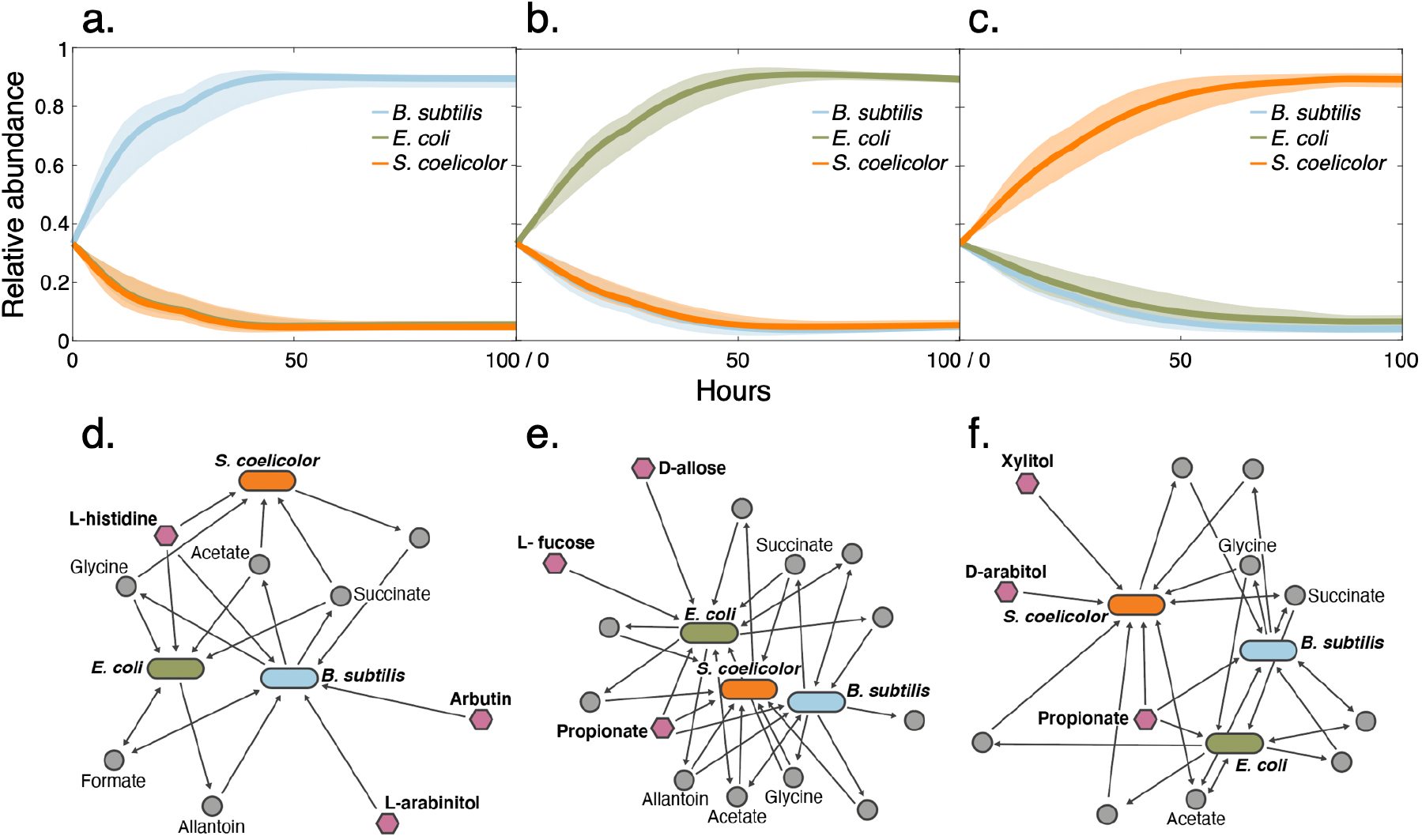
Simulated time-course trajectories of three-species community growth under various GA-determined environments. Genetic algorithm was used to determine environments that would allow *B. subtilis (a), E. coli (b*), and *S. coelicolor (c*) to reach abundances of 95% while the remaining organisms remained in the communities at basal levels. Dark lines indicate mean growth curves and shaded regions encompass the maximum and minimum relative abundances for each organism across 10 random environment seed sets. (*d-f*). Interaction network structures of representative environments that confer dominance to *B. subtilis (d*), *E. coli (e*), and *S. coelicolor (f*). Elongated ovals represent organisms, pink hexagons represent primary nutrients (environmental composition), and gray circles represent exchanged metabolites. Select commonly-exchanged metabolites are labeled.

We examined some of these environmental compositions in greater detail, identifying common interaction network structures that conferred the desired community phenotypes (Figure 4d-f). For example, in one of the environments that was selected to have *B. subtilis* dominate the community, our dFBA simulation revealed that it was the exclusive consumer of two out of three primary nutrients, while the third nutrient was shared between the three organisms (Figure 4d). A similar structure was also observed for the environments that optimized dominance of *E. coli* and *S. coelicolor* (Figure 4e, f), suggesting that nutrient specificity was a major driving force of organism dominance in these communities. We also observed dense networks of metabolic byproduct exchange, with molecules such as acetate, formate, glycine, and succinate being frequently transferred between organisms, paralleling previous experimental observations of organic acid transfer [67–69]. Given that a crucial element of our objective was for two organisms to persist at low abundances, these metabolic exchanges (along with consumption of a third primary nutrient) may be allowing the communities to remain stable with the desired taxonomic proportions.

## Discussion

The rational design of multispecies communities toward defined phenotypes is a challenging, yet enticing, goal of synthetic ecology. As the phenotypic traits of communities in complex settings remain difficult to predict [28,70,71], fulfilling this potential will require a synthesis of computational and experimental methods that focus on different aspects of these communities [10,72–74]. Here, we used *in silico* microbial communities to show how their ecological properties can be modulated via environmental modification, and presented a search algorithm to identify specific nutrient combinations that would result in desired phenotypes. We showed how this algorithm was quickly able to identify high-quality solutions for a variety of ecological objectives: from overall taxonomic balance to specific organism abundances and patterns of metabolic secretion and exchange. Given these capabilities, this method represents a computationally-inexpensive way to rapidly screen very large combinatorial spaces to produce desired community properties. Moreover, in addition to optimizing the various objectives tested here, our dFBA-genetic algorithm framework can be extended to encompass a greater number of important environmental attributes, such as varying nutrient concentrations and spatiotemporal nutrient variation [41,42].

Despite the flexibility and mechanistic insight afforded by a dFBA approach, engineering synthetic ecosystems *in vitro* will inevitably require experimental validation of modeling predictions. Our approach can be applied to this goal in two ways. First, *in silico* analogs of a desired experimental system may be iteratively screened as we have performed here, and the final environments generated by the genetic algorithm may then be explicitly tested experimentally. In this way, our method serves to generate an accessible number of testable hypotheses pertaining to specific ecological systems. Pairing of flux balance models and confirmatory experiments in this way has been used extensively to obtain a greater understanding of organism function as well as exploring previously unknown phenotypes [32,75–78]. However, as high-quality genome-scale reconstructions are limited to relatively few well-characterized model organisms, the applicability of this method is limited to a small set of community taxonomic compositions. A second strategy can forgo the dFBA component altogether, and use the evolutionary algorithm as a way to search through experimentally-derived community phenotypic data. As we showed how the GA was able to reach high-quality solutions with relatively few experimental data points, one could envision implementing a similar framework alongside the *in vitro* testing of a community. Here, iterative cycles of testing could be fed into a GA structure, which could inform the next stage of experiments. Given the increasing accessibility of high-throughput platforms for microbial ecology (e.g. microfluidics, microdroplets, etc.) [37–39], a search algorithm like ours can feasibly be deployed alongside such techniques to rapidly reach predefined community objectives.

## Methods

### Generation of environment-phenotype mapping with dFBA

We employed a dynamic flux balance analysis (dFBA) method [40] to test the response of a multispecies community in a combinatorial assortment of environments. This process, which was carried out using the COMETS (Computation of Microbial Ecosystems in Time and Space) software package [41,42], allowed us to extract a wide array of phenotypic data from simulated microbial communities. The process by which COMETS carries out these simulations has been outlined in detail in previous publications [33,41,42], and was carried out in the following way for our application: (1) Combinatorial environments were generated by combining an *in silico* minimal medium with limiting quantities of a set of carbon sources. This minimal medium, modeled after the composition of M9, contained molecules necessary for growth such as water, ions, and sources of nitrogen, phosphorus, and sulfur introduced at nonlimiting concentrations. Limiting amounts of 20 carbon sources were then added on an environment-by environment basis. These nutrients, an assortment of sugars, organic acids, and amino acids (Supplementary Table 2), were added in all combinations of up to 4 at equimolar ratios such that the total concentration of carbon in each environment was 50 mM C in 400 μL. This scheme resulted in 6,196 unique environmental compositions. (2) Genome-scale reconstructions [31,32] of 13 specific microbial organisms were placed in our *in silico* media compositions. These organism-specific models span a wide range of taxa and metabolic strategies, and were selected to maximize variation in endpoint community composition and interactions across our combinatorial environments (Supplementary Table 1). Based on an approximate total inoculum of OD600 0.05 corresponding to 1.6 × 10^7^ cells in 400 μL, and a cell mass of 2.8 × 10^−13^ grams dry weight (gDW) [79], all 13 organisms were inoculated into our *in silico* media at equal ratios of 3.45 × 10^−7^ gDW for a total inoculum of 4.48 × 10^−6^ gDW (OD600 0.05 total). The growth of these mixed cultures was then simulated in COMETS over the course of 24 hours, with a death rate parameter of 0.1 and a timestep of 0.01 hours [41]. Once completed, the total final biomass quantities, relative abundances, and secreted and absorbed metabolites for each environment were recorded.

For our second, exploratory application of the genetic algorithm, a larger pool of 154 carbon sources was used from which a maximum of three nutrients were selected per environment, resulting in 596,904 unique environmental compositions. Three organism genome-scale reconstructions (*B. subtilis, E. coli*, and *S. coelicolor* (Supplementary Table 1)) from our list of 13 were used and inoculated into our environments in a similar way to that described above. However, we did not explicitly simulate the community phenotypes in all combinatorial environments. Instead, only the environmental compositions selected by the genetic algorithm in each generation were tested and their performance recorded as above.

### Design and parametrization of genetic algorithm

A genetic algorithm is a search heuristic based on the principle of evolution by natural selection, which optimizes a particular objective function via the modification of a population of individual solutions [50]. Our selection of a genetic algorithm was based on its applicability to the optimization of nonlinear problems, which reflect the nature of complex environment-phenotype relationships in microbial communities. In our implementation, the individual solutions being modified are unique environmental compositions expressed as vectors denoting the presence of a particular nutrient. The objective function varied according to the phenotype being optimized. In this work, we selected a number of different objective functions to maximize, namely: (1) the overall Shannon entropy of a community as a reflection of taxonomic balance, (2) the relative abundances of each of the 13 *in silico* organisms, (3) the total number of metabolic exchanges, (4) the total metabolic flux directed at each of the 13 *in silico* organisms, (5) the total secretion flux of 24 different metabolic byproducts, and (6) the approximation of target relative abundances. The modifications of different solutions take place over the course of multiple ‘generations,’ in which each solution is scored according to the phenotype being optimized, and the best solutions are used to seed a new generation of candidate solutions. This process continues with the intent of converging on a set of optimal solutions.

Our implementation of the GA begins with a randomly-generated population made up of *P* environmental compositions. In order to demonstrate its extensibility to be used in parallel to an *in vitro* experimental system, we sought to minimize the number of environmental compositions *P* tested in each generation. Therefore, we limited the number of compositions to an experimentally-tractable *P* = 10 in each generation. The *P* environments were initialized with random assortments of up to *N* nutrients (N = 4 for our initial benchmarking study, and *N* = 3 for the second exploratory example). The community phenotypes resulting from each environment in the population (either pre-generated dFBA data in our benchmarking study, dFBA data generated as-needed in our exploratory example, and, in principle, experimental data if being used alongside an *in vitro* system) are recorded and used to rank each environment according to the objective function. The algorithm then selects the top *σ* environments to serve as ‘parents’ to the next generation of solutions. Having selected a set of *σ* parent environments, the algorithm then uses them to populate a new generation of *P* candidate solutions. This step takes place through processes of crossover (the individual elements of the parents are combined) and mutation (random, new elements are introduced). In our implementation, the parent nutrient vectors are linearized, and the remaining *P* – *σ* environments are populated with random assortments of the nutrients contained in the parent vector. Mutation then occurs, in which the individual nutrients of all *P* environments are subject to being randomly replaced by a nutrient yet unused in the set. The number of environments subject to crossover, as well as the probability of any individual nutrient being subject to mutation, are defined by crossover and mutation probabilities *p_c_* and *p_M_*, respectively.

We determined optimal values for the crossover and mutation probabilities *p_c_* and *p_M_* via a parameter grid search. To do this, we selected three representative objective functions: (1) maximization of community Shannon entropy, (2) maximization of the relative abundance of *B. subtilis*, and (3) maximization of the total number of metabolic exchanges. We then varied the values of *p_c_* from 0 to 1 in intervals of 0.1, and the values of *p_M_* from 0 to 0.45 in intervals of 0.05. The values of *p_M_* were maintained under 0.5 in order to ensure the GA process would not diverge from optimal solutions via excessive mutation. For each pairing of *p_c_* and *p_M_*, we applied our GA 50 times, each with a random seed set of *P* = 10 different environments. We then evaluated the performance of the GA for each objective using a performance score S. This score is based on a combination of two metrics: (1) the number of generations required for a set of solutions to surpass the 99^th^ percentile of a given objective (G_99_) and (2) the percentile reached at the final generation of the algorithm *Pr_end_*. Since a lower *GAA* denotes better performance, the performance score S is defined as follows:

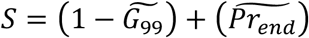

Where 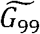 and 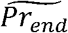 are normalized from 0 to 1, such that S can range from 0 to 2. We found that the best [*p_c_,p_M_*] values were [0.7, 0.45] for our first objective, [1, 0.3] for our second, and [0.8, 0.45] for our third (Supplementary Figure 2). Interestingly, while our *p_c_* values were consistent with commonly-used crossover parameter values [80], we noticed low variability in performance scores S with respect to changing mutation probabilities *p_M_.* We thus used an average best [*p_c_, p_M_*] values ([0.8, 0.4]) for all of our GA objectives. For a population size of *P =* 10, a *p_c_* value of 0.8 therefore results in *σ* = 2 environments being selected to carry over and seed a new generation. Once a new generation of environments is created via carryover, crossover, and mutation, the community phenotype in in each environment is recorded, and the environments are ranked. In our implementation, this process of selection continues until a maximum of generations *G* is reached (Figure 2).

## Data accessibility

MATLAB scripts for running the genetic algorithm are available at github. com/segrelab/EvolutionaryAlgorithms.

## Authors’ contributions

A.R.P and D.S. designed the research. A.R.P. developed the algorithm framework, collected data, and designed and performed simulations. A.R.P. and D.S. wrote the manuscript. Both authors read and approved the final manuscript.

## Competing interests

The authors declare that no competing interests exist in relation to this manuscript.

## Funding

A.R.P. is supported by a Howard Hughes Medical Institute Gilliam Fellowship and a National Academies of Sciences, Engineering, and Medicine Ford Foundation Predoctoral Fellowship. We gratefully acknowledge support from the U.S. Department of Energy, Office of Science, Office of Biological & Environmental Research through the Microbial Community Analysis and Functional

Evaluation in Soils SFA Program (m-CAFEs) under contract number DE-AC02-05CH11231 to Lawrence Berkeley National Laboratory, as well as the National Institutes of Health (grants 5R01DE024468, R01GM121950), the National Science Foundation (grants 1457695 and NSFOCE-BSF 1635070), the Human Frontiers Science Program (grant RGP0020/2016), and the Boston University Interdisciplinary Biomedical Research Office.

## Acknowledgements

The authors wish to thank members of the Segrè lab for inspiring conversations. We are especially grateful to Joshua Goldford, Mark Kon, and Dileep Kishore for helpful discussions, and to David Bernstein, Melisa Osborne, and Devlin Moyer for their constructive comments on the manuscript.

## Notes

### Competing Interest Statement

The authors have declared no competing interest.

