## Supplementary Materials for "An evolutionary algorithm for designing microbial communities via environmental modification"

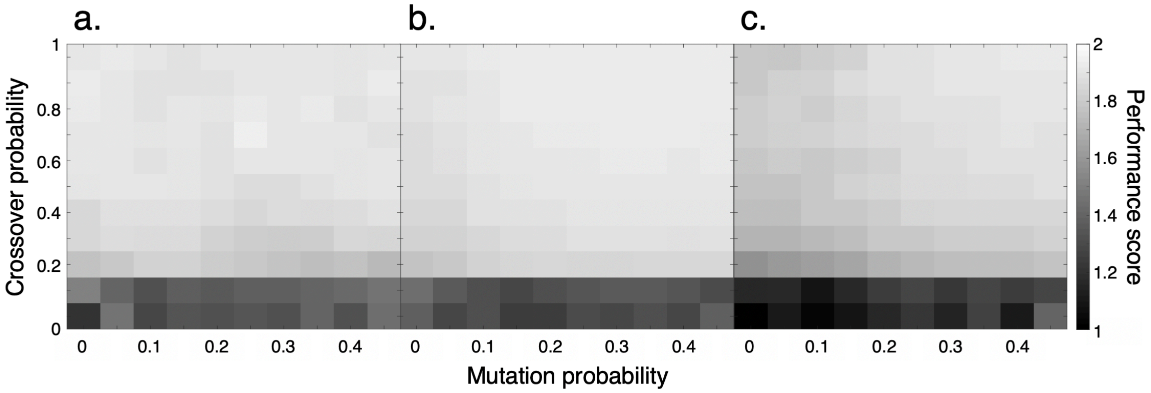


Supplementary Figure 1. Genetic algorithm parameter grid search results. The crossover probability $p_C$ and mutation probability $p_M$ of the algorithm were varied, and a performance score was calculated for each pair based on (1) the number of generations required for the algorithm to reach the 99$^{th}$ percentile of a given objective and (2) the percentile of the solution at the last generation (see Methods). Three representative ecological phenotypes were selected to perform the grid search: *(a)* maximization of community Shannon entropy, *(b)* maximization of the relative abundance of *B. subtilis*, and *(c)* maximization of the total number of metabolic exchanges. For each $[p_C, p_M]$ parameter pairing, the mean of 50 random environment seed sets is shown. Using an additional moving average smoothing procedure, this search process resulted in generally consistent parameter values emerging (the best $[p_C, p_M]$ pairings were identified as [0.7, 0.25] for *(a)*, [1, 0.45] for *(b)*, and [1, 0.4] for *(c)*), which informed the decision to use an average of these values ([0.9, 0.35]) for all search processes.

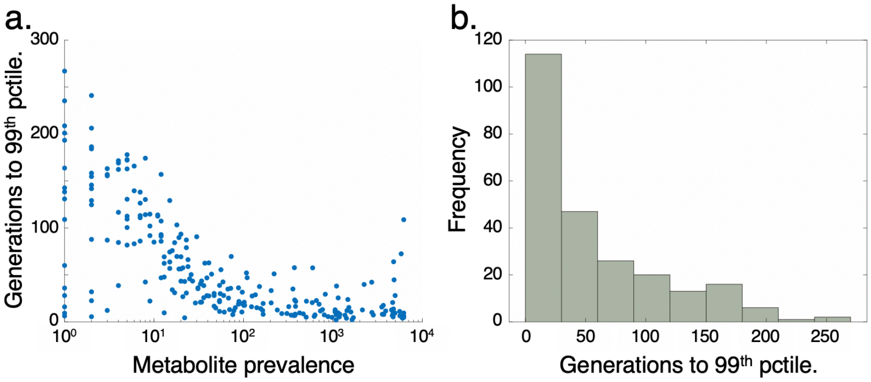


Supplementary Figure 2. GA performance according to prevalence of secreted metabolites. *(a)*. Mean number of generations required for GA to find environments that surpass the 99$^{th}$ percentile of total metabolite secretion flux ($n = 246$ organic secreted metabolites). For each data point, the mean of 10 random environment seed sets is shown. *(b)*. Frequency of required generations to reach aforementioned objective. Environments that maximize the secretion of a majority of metabolites (61.4%) can be identified within 50 generations.

Supplementary Table 1. Genome-scale models of organisms used in simulations of community growth.

| **Organism name** | **Reference** |
|---|---|
| *Bacillus subtilis* | [1] |
| *Escherichia coli* | [2] |
| *Klebsiella pneumoniae* | [3] |
| *Lactococcus lactis* | [4] |
| *Pseudomonas aeruginosa* | [5] |
| *Porphyromonas gingivalis* | [6] |
| *Rhodobacter sphaeroides* | [7] |
| *Shigella boydii* | [8] |
| *Saccharomyces cerevisiae* | [9] |
| *Streptomyces coelicolor* | [10] |
| *Salmonella enterica* | [11] |
| *Shewanella oneidensis* | [12] |
| *Zymomonas mobilis* | [13] |

Supplementary Table 2. List of 20 limiting carbon sources for initial dFBA environment-phenotype mapping.

| **Nutrient name** |
|---|
| Acetate |
| 2-Oxoglutarate |
| Butyrate |
| Citrate |
| D-Fructose 6-phosphate |
| L-Cysteine |
| Fumarate |
| D-Glucose 6-phosphate |
| L-Isoleucine |
| D-Glucose |
| Glycerol |
| D-Lactate |
| L-Histidine |
| D-Malate |
| L-Arginine |
| L-Phenyalanine |
| Pyruvate |
| Succinate |
| Sucrose |
| Trehalose |

Supplementary Table 3. Mean number of generations (across 50 random seed environments) required for genetic algorithm to exceed the 99th percentile of a given objective ($G_{99}$).

| Objective | | | $G_{99}$ |
|---|---|---|---|
| Overall Shannon Entropy | | | 3.16 ± 0.49 |
| Total number of total metabolic exchanges | | | 8.12 ± 0.86 |
| Maximization of organism relative abundances | *B. subtilis* | | 6.12 ± 0.57 |
| | *E. coli* | | 4.02 ± 0.49 |
| | *K. pneumoniae* | | N/A |
| | *L. lactis* | | N/A |
| | *P. aeruginosa* | | 3.24 ± 0.60 |
| | *P. gingivalis* | | N/A |
| | *R. sphaeroides* | | N/A |
| | *S. boydii* | | 13.10 ± 2.61 |
| | *S. cerevisiae* | | N/A |
| | *S. coelicolor* | | 3.26 ± 0.46 |
| | *S. enterica* | | 9.06 ± 1.20 |
| | *S. oneidensis* | | 2.08 ± 0.24 * |
| | *Z. mobilis* | | N/A |
| Number of metabolic exchanges toward organisms | *B. subtilis* | | 9.44 ± 1.16 |
| | *E. coli* | | 12.92 ± 2.50 |
| | *K. pneumoniae* | | N/A |
| | *L. lactis* | | N/A |
| | *P. aeruginosa* | | 12.8 ± 2.43 |
| | *P. gingivalis* | | N/A |
| | *R. sphaeroides* | | N/A |
| | *S. boydii* | | 12.16 ± 1.75 |
| | *S. cerevisiae* | | N/A |
| | *S. coelicolor* | | 9.46 ± 0.89 |
| | *S. enterica* | | 11.22 ± 1.27 |
| | *S. oneidensis* | | 9.78 ± 0.97 |
| | *Z. mobilis* | | N/A |
| Total metabolite secretion flux | Most-secreted metabolites | Acetate | 5.78 ± 0.45 |
| | | Formate | 6.90 ± 0.55 |
| | | Ethanol | 7.16 ± 0.70 |
| | | Succinate | 7.74 ± 0.78 |
| | | Glycine betaine | 33.91 ± 4.25 |
| | | Glycine | 13.06 ± 2.10 |
| | | Oxaloacetate | 9.44 ± 1.27 |
| | | Isocitrate | 7.28 ± 0.76 |
| | | Phenylacetate | 7.04 ± 0.88 |
| | | Propionate | 11.46 ± 1.58 |
| | | L-malate | 11.76 ± 1.48 |

| | | | |
|---|---|---|---|
| | | L-alanine | 11.52 ± 1.14 |
| | Least-secreted metabolites | Spermidine | 173.56 ± 11.48 |
| | | L-rhamnose | 74.20 ± 9.94 |
| | | Deoxyribose | 100.27 ± 11.17 |
| | | D-allose | 184.47 ± 10.55 |
| | | 4-hydroxy-L-threonine | 173.27 ± 9.08 |
| | | Hexadecanoate | 159.95 ± 11.19 |
| | | 3-methylbutanal | 156.32 ± 10.06 |
| | | L-ascorbate | 139.68 ± 11.46 |
| | | D-carnitine | 155.12 ± 11.52 |
| | | L-idonate | 197.76 ± 10.77 |
| | | D-glucuronate-1-phosphate | 87.75 ± 12.02 |
| | | Thymidine | 148.13 ± 11.53 |

* For *S. oneidensis*, the 99th and 100th percentiles of relative abundances both equaled 1 in our dataset, and as such the generation at which the 99th was passed is undefined. The quantity shown is therefore the generation at which the 98th percentile was passed.

Supplementary Table 4. List of most and least highly-secreted metabolites across all dFBA simulations.

| **Metabolite** | **Secretion flux** |
|---|---|
| Most secreted metabolites | |
| Acetate | 1.09E+06 |
| Formate | 8.63E+05 |
| Ethanol | 1.29E+05 |
| Succinate | 1.04E+05 |
| Glycine betaine | 6.19E+04 |
| Glycine | 5.84E+04 |
| Oxaloacetate | 2.96E+04 |
| Isocitrate | 2.33E+04 |
| Phenylacetate | 1.22E+04 |
| Propionate | 1.14E+04 |
| L-malate | 1.13E+04 |
| L-alanine | 8.94E+03 |
| Least secreted metabolites | |
| Spermidine | 7.12E-12 |
| L-rhamnose | 6.55E-12 |
| Deoxyribose | 5.66E-12 |
| D-allose | 5.62E-12 |
| 4-hydroxy-L-threonine | 4.44E-12 |
| Hexadecanoate | 3.73E-12 |
| 3-methylbutanal | 3.45E-12 |
| L-ascorbate | 2.97E-12 |
| D-carnitine | 2.22E-12 |
| L-idonate | 2.06E-12 |
| D-glucuronate-1-phosphate | 1.69E-12 |
| Thymidine | 1.04E-12 |
